# Bridging the Gap: From Neuroanatomical Knowledge to Tractography of Brain Pathways

**DOI:** 10.1101/2020.08.01.232116

**Authors:** Guillermo Gallardo, Demian Wassermann, Alfred Anwander

## Abstract

Despite recent advances in tractography, the gap remains wide between the descriptions of white-matter pathways in the literature and the methods to reconstruct and study them from dMRI images. Here, we tackle this challenge by proposing a language to define white matter tracts, namely WMQL^T^, and a tool to automatically reconstruct pathways from their WMQL^T^ queries. Our method is performant, flexible enough to allow defining tracts using multiple modalities, and allows to extend ROI-based reconstruction methods. Leveraging our language, we define 19 major brain tracts, alongside their subdivisions, and reconstruct them in a large population. We show that the shape of the reconstructed pathways, as well as their connectivity and lateralizations are in accordance with the current neuroanatomical literature. Finally, we showcase our technique in two scenarios: computing the functional subdivisions of a tract, and assessing the role of handedness and gender in the lateralization of language-related tracts.

## 1. Introduction

Diffusion tractography enables reconstructing white matter pathways non invasively. Since manually dissecting brain pathways is a time consuming and error prone process, different automatic reconstruction methods have been recently developed (Zhang *et al*., 2008; Yendiki *et al*., 2011; Yeatman *et al*., 2012; Warrington *et al*., 2020). By predefining regions and rules to guide the tractography process, automatic region-based methods facilitate the systematic study of tracts in large populations. However, extending such methods beyond their predefined set of tracts remains challenging for the non-expert users. This is due to the inherent difficulty in manually translating anatomical descriptions from the literature to method-specific regions and rules.

Current region-based methods reconstruct white-matter pathways by defining rules and regions of interest (ROIs) for tractography. The ROIs are generally delimited in a standardized space, while the rules define which ROIs a tract must traverse, avoid or end in (Mori and van Zijl, 2002). Based on this approach Wakana et al. (2007) presented recipes to manually locate ROIs for 11 major bundles. Wakana’s recipes combine anatomical landmarks with color coded fractional anisotropy (FA) maps, in order to guide the manual placement of either one or two rois per tract. Catani et al. (2008) presented a similar approach, combining positions in the MNI space with color coded FA maps in order to define recipes for 10 major bundles. Yeatman et al. (2012) took a step further with their tool AFQ. AFQ isolates tracts by warping the Wakana’s ROIs (Wakana et al. 2007) into the individual MRI space, and uses them to filter a whole-brain tractography. Using a similar approach, Warrington et al. (2020) recently introduced XTRACT, a tool that automatically dissects 22 major tracts in both humans and macaques. XTRACT works by warping ROIs from a template space (MNI for humans, F99 for macaque’s) to the MRI space of the subject, and using them to perform tractography using FSL (Jenkinson *et al*., 2012). Finally, extending the ROI placement methods are the bundle specific tractography methods (Yendiki et al. 2011; Rheault et al. 2019; Wasserthal et al. 2019). These methods add orientation priors to the tractography process, guiding it through regions with poor directionality or high amount of crossing.

Reconstructing tracts through ROI placement methods provides accuracies comparable to that of manual dissections, while achieving high reproducibility across subjects (Wakana *et al*., 2007; Catani and Thiebaut de Schotten, 2008; Warrington *et al*., 2020). However, the methods dependency on volumetric registrations impose limitations in their use case scenarios. By warping binary ROIs from anatomical templates, current tools preclude using regions that do not closely follow anatomical landmarks or present high spatial variability (Coalson, Van Essen and Glasser, 2018). Furthermore, extending a method beyond their predefined set of tracts requires the user to manually delimit ROIs in the method’s template, relying on their technical knowledge. To tackle these issues, Wassermann et al. (2016) introduced the white matter query language (WMQL). The WMQL formalizes tract descriptions in a human readable computer language, allowing to create and modify definitions without the need of an engineering background. Moreover, WMQL definitions are not tied to anatomical constraints, since regions from any brain atlas can be used to characterize a tract. Alongside the language, Wasserman et al. (2016) presented a tool to extract 14 major tracts, and their subdivisions, from a set of streamlines. However, the tool requires the user to pre-compute a whole-brain tractography for every subject. This translates into an inefficient process, where millions of streamlines are computed in order to filter a few representatives per tract.

In this work, we build on WMQL and ROI methods to present a novel technique to easily define and automatically reconstruct pathways in the brain. Our technique can be used by itself, or to extend current ROI methods. Specifically, we (i) simplify WMQL’s syntax, to create a language better suited for creating tracking masks that we name WMQL^T^; (ii) refine existing WMQL definitions; (iii) develop an interpreter that translates WMQL^T^ into subject specific tracking masks; and (iv) implement a tractography pipeline to reconstruct the desired tracts. Using WMQL as a base allows our tract definition, and therefore our reconstruction pipeline, to be easily extended by any user. However, WMQL^T^ presents many advantages with respect to WMQL. First, our method can define and reconstruct tracts using multiple atlases at the same time. This enables to define tracts using multiple modalities, i.e. by using an anatomical and functional atlas concurrently. Furthermore, by creating tracking masks we gain performance and precision through a controlled seeding with respect to WMQL. Finally, a major advantage of our technique is that our tracking masks can be used to extend existing reconstruction methods, giving the user freedom to use their prefered framework.

We validate our method in 600 subjects from the Human Connectome Project (HCP) dataset (Van Essen *et al*., 2012). We show how our technique consistently reconstructs 19 major brain tracts correctly, and we derive: (i) a probabilistic atlas of white matter tracts, and (ii) a probabilistic atlas of brain connectivity. Also, to showcase how our interpreter can aggregate information from different atlases, we reconstruct the motor-related functional subdivisions of the corticospinal pathway. We further validate our results by computing the lateralization of language-related tracts and showing that our results are in agreement with current neuroanatomical knowledge. Finally, we apply our technique to compare different lateralization indices used in the literature, and assess the role of handedness and gender in the lateralization of language-related tracts.

This paper is organized as follows: In the Methods section we present our formalization of WMQL^T^, alongside our interpreter. In the Experiments and Results section we present our results on the HCP data. We then discuss our results with respect to current neuroanatomical knowledge, and provide our conclusions.

## 2. Methods

### 2.1. The White Matter Query Language for Tracking

Based on WMQL (Wassermann et al., 2016), the white matter query language for tracking (WMQL^T^) formalizes the anatomical description of fiber pathways in a near-to-English textual computer language. In contrast with WMQL, which is intended to filter streamlines from a precomputed whole-brain tractogram, WMQL^T^ is designed to create tracking masks. These tracking masks can then be used in any tractography algorithm to reconstruct the desired tract. As in WMQL, WMQL^T^ describes tracts as relationships between regions of an atlas. Formally, the syntax of WMQL^T^ can be expressed using the Backus-Naur form for the description of context-free grammars as:

~~~
RELATIONS: = or | and
ROI: = atlas-roi | atlas-roi RELATIONS ROI |
       anterior_of(ROI) | posterior_of(ROI) |
       medial_of(ROI) | lateral_of(ROI) |
       superior_of(ROI) | inferior_of(ROI)
TRACT:= ROI | not_in(ROI) | endpoints_in(ROI) | TRACT and TRACT
~~~

A tract in WMQL^T^ syntax is constructed by specifying regions of interest (ROIs), and rules on how to use them in tractography. The ROIs are defined as: (1) regions from an atlas (atlas-roi); (2) relative positions with respect to atlas regions (i.e. anterior_of, posterior_of); and (3) logical operations between ROIs. The rules meanwhile define which regions the tract must avoid (not_in) or traverse (ROI, endpoints_in). It’s important to note that, in order to maintain the language simple, we kept the term endpoints_in from WMQL. However, while current tractography algorithms allow to denote that a region must be traversed (known as inclusion, or waypoint masks), avoided (exclusion masks), or connected to (seed masks), it is not possible to indicate that the tract must finish in a region. Therefore, in WMQL^T^ endpoints_in refers to the ROIs that the tract must connect, without necessarily ending in.

The expressive power of WMQL^T^ allows to define white matter tracts in a simple yet unequivocal way. For example, take the following definition of the Inferior Fronto-Occipital Fascicle (IFOF) (Catani and Thiebaut de Schotten, 2008): “*The inferior fronto-occipital fasciculus is a ventral associative bundle that connects the ventral occipital lobe and the orbitofrontal cortex. […]*”. Using the Desikan atlas (Desikan et al., 2006) as implemented in freesurfer, we can translate this definition to WMQL^T^:

~~~
Cuneus.left: = 1005
Precentral.left: = 1024
Insula.left = 1035
Superior_Frontal.side:= 1028
White_Matter.left:= 41
…
Frontal.left:= IFG.left or Middle_Frontal.left or
               Superior_Frontal.left or Orbito_Frontal.left
Occipital.left:= Cuneus.left or Lateral_Occipital.left or
                 Lingual.left or Pericalcarine.left
Ventral_ROI.left:= White_Matter.left and inferior_of(Precentral.left)
Dorsal_ROI.side:= White_Matter.side and superior_of(Insula.side)
IFOF.left:= endpoints_in(Occipital.left) *and*
            endpoints_in(Frontal.left) and
            Ventral_ROI.left and
            not in Dorsal_ROI.side
~~~

As in WMQL, WMQL^T^ also includes the suffix “.side”, to help reuse a definition in both hemispheres, and the “import” command, which allows to load an existing definitions file. In this way, the previous IFOF definition can be drastically simplified by predefining all the freesurfer definitions in a file.

~~~
import Freesurfer.qry
Dorsal_ROI.side = White_Matter.side and superior_of(Insula.side)
Ventral_ROI.side = White_Matter.side and inferior_of(Precentral.side)
IFOF.side:= endpoints_in(Occipital.side) *and*
            endpoints_in(Frontal.side) and
            Ventral_ROI.side and
            not in Dorsal_ROI.side
~~~

This IFOF definition highlights how easy it is to both define and interpret white matter pathways written in WMQL^T^. Furthermore, it’s important to remark that WMQL^T^ does not depend on a particular atlas, allowing to create definitions using functional localizers, or anatomical landmarks based on individualized landmarks.

### 2.2. Interpreting WMQL^T^: Translating Definitions into Tracking Masks

To translate WMQL^T^ definitions into tracking masks, we implemented a parser. Our parser takes as input a definition file, and a set of atlases. In contrast to the WMQL parser (Wasserman et al. 2016), which takes as input a single atlas, ours allows to combine different modalities in the definition of the pathway (i.e. function and structure). Our parser implements a left-to-right parsing algorithm to translate WMQL^T^ definitions into an abstract tree. This tree is then transversed and translated into ROI masks for tractography. In order to transform WMQL^T^ definitions into tracking masks, we evaluated relations, terms and clausules into operations on binary masks. In our implementation, we keep in memory 3 sets of masks: *seed, traverse*, and *avoid*. Then, the WMQL^T^ operations are evaluated as follows: (i) Each region named in the definition is extracted from the corresponding atlas as a binary mask. (ii) The boolean operations *“or”, and “and”* are computed as voxelwise operations over masks. (iii) Each relative positional clause is transformed into a binary mask by setting to 1 all the voxels in the correspondent anatomical direction. For example, the clause anterior_of(amygdala) will return a binary image where each voxel anterior to the amygdala is set to 1. (iv) The term *endpoints_in(A)* defines A as an *end* mask. (v) The term *not_in B* defines B as an *avoid* mask. (vi) Finally, every mask not assigned as *end* or *avoid*, is stored as a *traverse* mask. For example, for the IFOF definition given in the previous section, our parser will create one *traverse* mask, one avoid mask, and two end masks (see Figure 1).

**Figure 1.**
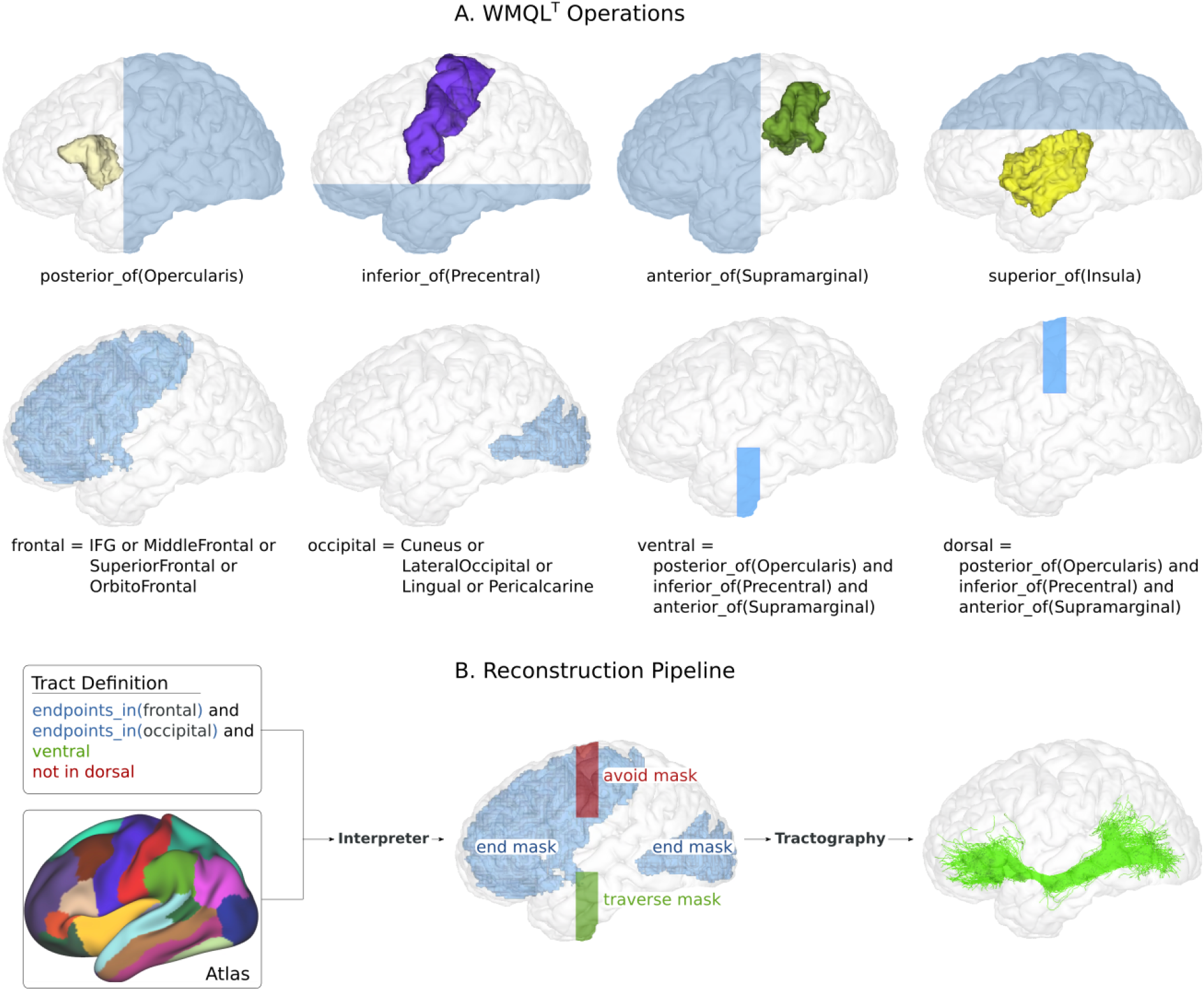
A. Examples on how different WMQL^T^ terms are interpreted and transformed into tracking masks. B. Overview of our reconstruction pipeline. Our interpreter translates WMQL^T^ into end, avoid, and traverse masks. In order to reconstruct the desired tract, our tractography pipeline uses the end and traverse masks as seeding and inclusion, and the avoid mask as exclusion.

### 2.3. Reconstructing the Tracts: From WMQL^T^ to Probabilistic Tractography

Our interpreter translates WMQL^T^ definitions into binary masks denoting the regions the tract is expected to traverse, and avoid. These masks can be readily used to inform any tractography algorithm (Jenkinson *et al*., 2012; Garyfallidis *et al*., 2014; Tournier *et al*., 2019). A simple reconstruction would require using the *avoid* masks for exclusion, the *traverse* masks for seeding, and the *end* masks for inclusion (sometimes called waypoints in the literature). In this way, the algorithm tracks from the tract’s core, and uses the end-voxels as inclusion-points to ensure reconstructing the correct morphology (Warrington *et al*., 2020). Although more time consuming, a better approach is to also use the traverse masks for inclusion, and the end masks as seeding-points. This allows to further reconstruct the pathway from its extremes, better characterizing its overall connectivity.

The tracking masks produced by our interpreter can also be used to extend current tract-specific reconstruction methods. Using as input a template-based volume atlases, as those defined in the MNI (humans) or F99 (macaque monkey) space, our parser will create tracking-masks in such template space. The resulting masks are compatible with current reconstruction methods (i.e. XTRACT, AFQ), allowing the user to easily extend them beyond their predefined set of tracts.

### 2.4. A Pipeline for WMQL^T^-Informed Probabilistic Tractography

To ease the process of reconstructing subject-specific tracts, we developed a pipeline that automatizes translating WMQL^T^ queries into brain pathways. Our pipeline takes as input a set of WMQL^T^ definitions, one or more subject-specific atlas, and a diffusion image. We first estimate the diffusion Fiber Orientation Distributions (FODs) (Tournier et al., 2004) from the Constrained Spherical Deconvolution (CSD) model (Tournier et al., 2007) implemented in MRtrix3 (Tournier *et al*., 2019). Then, we use our WMQL^T^ interpreter to obtain tracking masks from the individual segmentation and parcelation provided by e.g. freesurfer. We use the avoid masks for exclusion, and the traverse masks for both seeding and inclusion. Next, we reduce the end masks to the voxels at the white-gray matter interface, and use them as both seeding and inclusion masks in the default MRtrix probabilistic tracking algorithm. We simulate 8 streamlines per seed-voxel, constraining the tracking to the white matter volume of the subject.

### 2.5. Refining Previous WMQL Definitions

We refined the WMQL definition of 19 major tracts (Wassermann et al. 2016) to better suit them to the tractography process. Particularly, we redefined the arcuate fascicle (AF); cingulum bundle (CB); corpus callosum (I-VII); cortico-spinal tract (CST); inferior fronto occipital fascicle (IFOF); inferior longitudinal fascicle (ILF); middle longitudinal fascicle (MLF); superior longitudinal fascicle (SLF I, II, III); uncinate fascicle (UF); and connections to the thalamus and striatum. Our refined definitions, alongside the type and amount of masks created per tract, can be found in Table 3. A consistent change across tracts is the addition of exclusion regions. This allows to further constrain the shape of the tracts, thus reducing the amount of spurious streamlines created.

## 3. Experiments and Results

In the previous section we introduced a method to automatically reconstruct white matter pathways from human readable descriptions of neuroanatomy. Now, we validate our technique by showing how it correctly reconstructs 19 major tracts in 600 subjects. For this, we derive for each tract a population-wise map of white-matter location and cortico-cortical connectivity. Such maps denote in which percentage of the population a tract is present in a specific location, or connects to it, naturally shielding probabilistic maps of position and connection per tract. We compare the maps of each tract with their literature counterpart. Also, to showcase how our method can define tracts using multi-modal information, we reconstruct the motor-related functional subdivisions of the corticospinal pathway. To further validate our technique, we compute the lateralization of language-related pathways. We show that our results are consistent with those thoughtfully documented in the literature (Catani and Thiebaut de Schotten, 2008; Thiebaut de Schotten, *et al*., 2011; Thiebaut de Schotten, *et al*., 2011; Warrington *et al*., 2020). Finally, after validating our technique, as an application example of our method, we use it to assess current theories of handedness and gender influence on the lateralization of the language-related pathways (Thiebaut de Schotten, *et al*., 2011; Gierhan, 2013; Howells *et al*., 2018).

### 3.1. Data and Preprocessing

A total of 600 subjects (300 males and 300 females, 75 left-handed, ages 20-60) were randomly selected from the group S1200 of the Human Connectome Project (HCP). In this project, the acquisition protocol was optimized for high spatial resolution and signal-to-noise ratio. The acquisition included two datasets with LR and RL polarity of the EPI phase encoding direction: TE/TR 89ms/5.5s; 1.25mm isotropic voxels; b = 0 (6 images), and 271 directions equally distributed across 3 b-shells of 1000, 2000, and 3000s/mm^2^. We refer the interested reader to (Sotiropoulos et al., 2013) for further details of the acquisition and preprocessing pipeline. Every subject has been already preprocessed with the HCP minimum pipeline (Glasser et al., 2013), for which they possess ready to use: diffusion image, T1 weighted image, and cortical surface reconstructions. The dataset provides a registration of all cortical surfaces to a common space which is labeled by different atlases, like the gyral based parcellation atlas (Desikan et al., 2006). Furthermore, each surface based parcellation is also available (or easily computable) in the volumetric voxel space.

### 3.2. Reconstructing White-matter Tracts

Using our tractography pipeline, we reconstructed 19 major tracts per hemisphere on each subject: Arcuate fascicle (AF), cingulum bundle (CB), uncinate fascicle (UF), superior longitudinal fascicle (SLF I, II, III), inferior fronto-occipital fascicle (IFOF), middle longitudinal fascicle (MLF), uncinate fascicle (UF), corpus callosum (CC I, II, III, IV, V, VI, VII), corticospinal tract (CST), Thalamus connections, and Striatal connections. Briefly, our pipeline: (i) translated the WMQL^T^ definitions (see table 3) into seeding, inclusion and exclusion masks; (ii) project the seeding masks, in order to track from the white-gray matter interface; and (iii) used the resulting masks to guide the tracking on a state-of-the-art tractography algorithm.

Figure 2 shows the streamlines reconstructed for each tract on a single subject. The figure shows how the obtained streamlines correctly reconstruct the general shape which was expected for each pathway.

**Figure 2.**
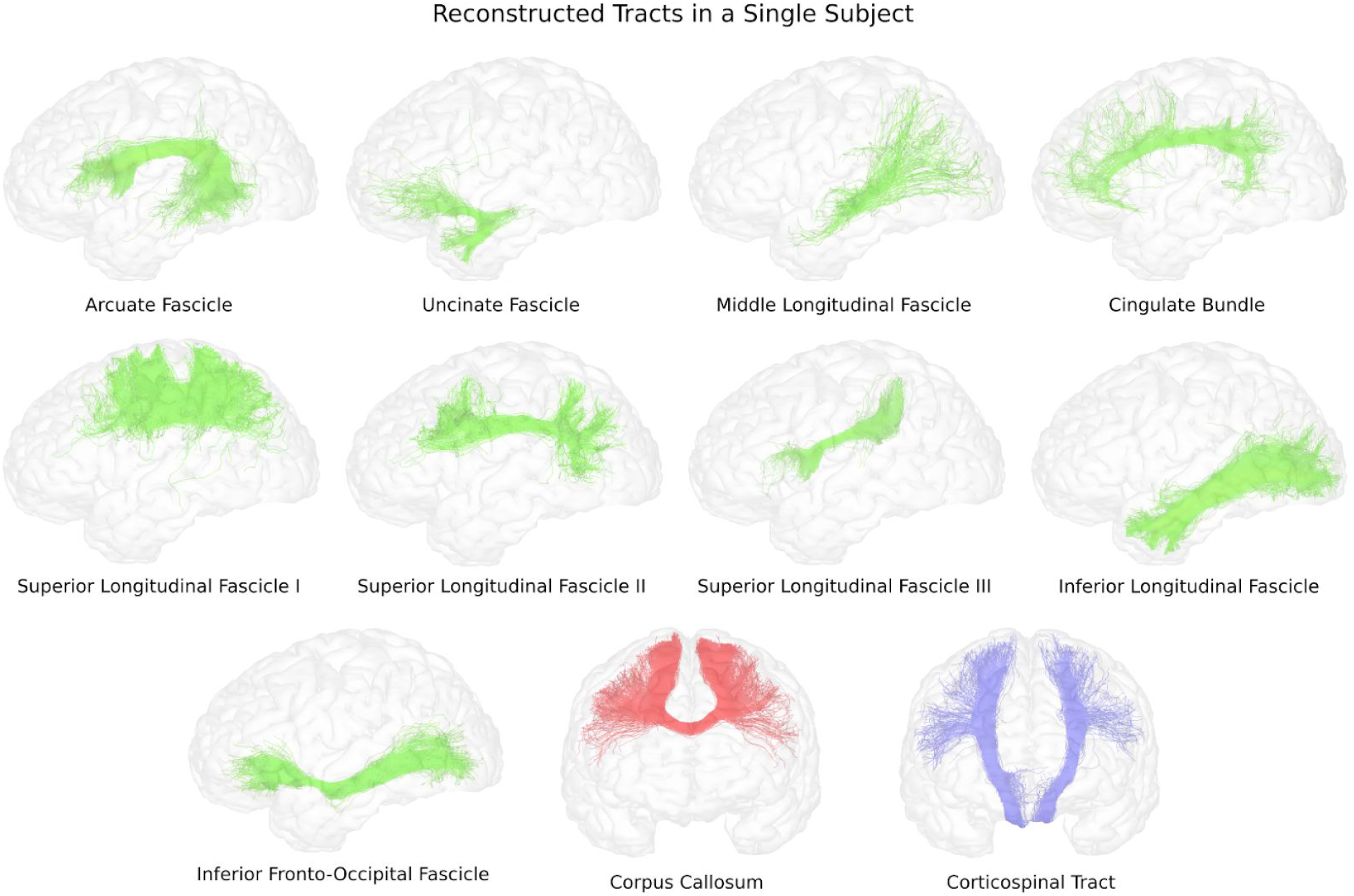
Examples of pathways reconstructed using our technique in a single subject from the Human Connectome Project.

### 3.3. Combining Atlases to Refine and Enrich Definitions

Our tool allows us to further combine different atlases in order to refine and enrich anatomical definitions of tracts. In order to showcase its possibilities, we reconstructed the motor-related functional subcomponents of the Corticospinal Tract (CST) of one subject. We derived a functional atlas from the average responses of 100 subjects to different stimuli (<u>Barch et al., 2013</u>), readily available in the S1200 release of the HCP. We thresholded the map with the z-scores related to the movement of the hand, foot and tongue in the right hemisphere using the 65% of their highest value. This resulted in 3 distinguishable regions across the precentral gyrus. With WMQL^T^ we defined 3 tracts, representing the functional subcomponents of CST. Here we exemplify one:

~~~
import Freesurfer.qry
foot.right:= 11
CST_foot.right:= endpoints_in(motor_cortex.right) and
               endpoints_in(foot.right)
~~~

Finally, we gave the definitions, alongside the Desikan atlas and the functional atlas to our WMQL^T^ interpreter. Figure 3 shows the reconstructed bundle, alongside the 3 functional subcomponents that we obtained.

**Figure 3.**
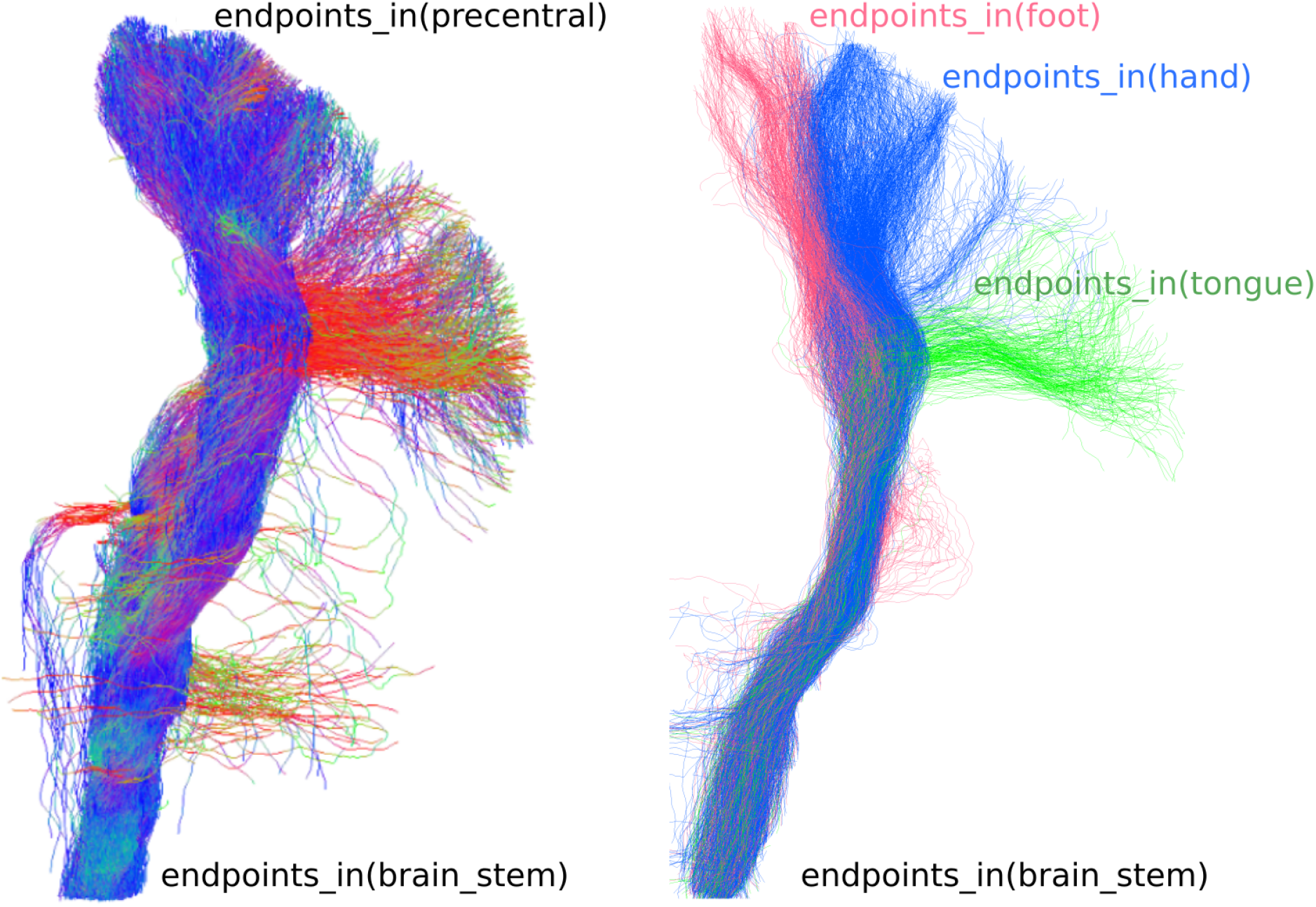
WMQL^T^ allows to define tracts by mixing different types of definitions, as anatomical and functional ones. Left. Corticospinal tract reconstructed using regions from an anatomical atlas (Desikan *et al*., 2006). Right. Functional subdivisions of the corticospinal tract, obtained by including functional information into the tract’s definition. The used functional atlas was derived from the subject’s response to motor-related functional stimuli (Barch *et al*., 2013).

### 3.4. Population Maps of Reconstructed White-matter Tracts

To assess the correct shape of each reconstructed tract at the population level, we transformed the streamlines of each subject into the MNI space, using the registrations already available in the HCP dataset. Once in the common space, we computed the binary visitation map of each tract across subjects (Thiebaut de Schotten *et al*., 2011). A visitation map shows how many streamlines intersect each voxel in the target space. In general, those visitation maps generated by probabilistic tractography must be thresholded to remove spurious false-positive pathway trajectories and endings before the visitation map can be analysed. This threshold is arbitrary and depends on the respective application. For the generation of population maps, we choose to binarize the individual visitation map using all voxels with at least one streamline. Finally, we aggregated the binary tract maps across subjects. This population average map naturally shields a probabilistic white matter atlas, which estimates the probability of finding each tract in a specific position.

Figure 4 shows a 3D reconstruction of the probabilistic maps derived after thresholding at the level of 10%. This is, the figure highlights the voxel in which at least 10% of the population had at least one streamline crossing it. The reconstructed volumes closely follow the expected core-body of each tract, showing that our method consistently reconstructs the tracts correctly.

**Figure 4.**
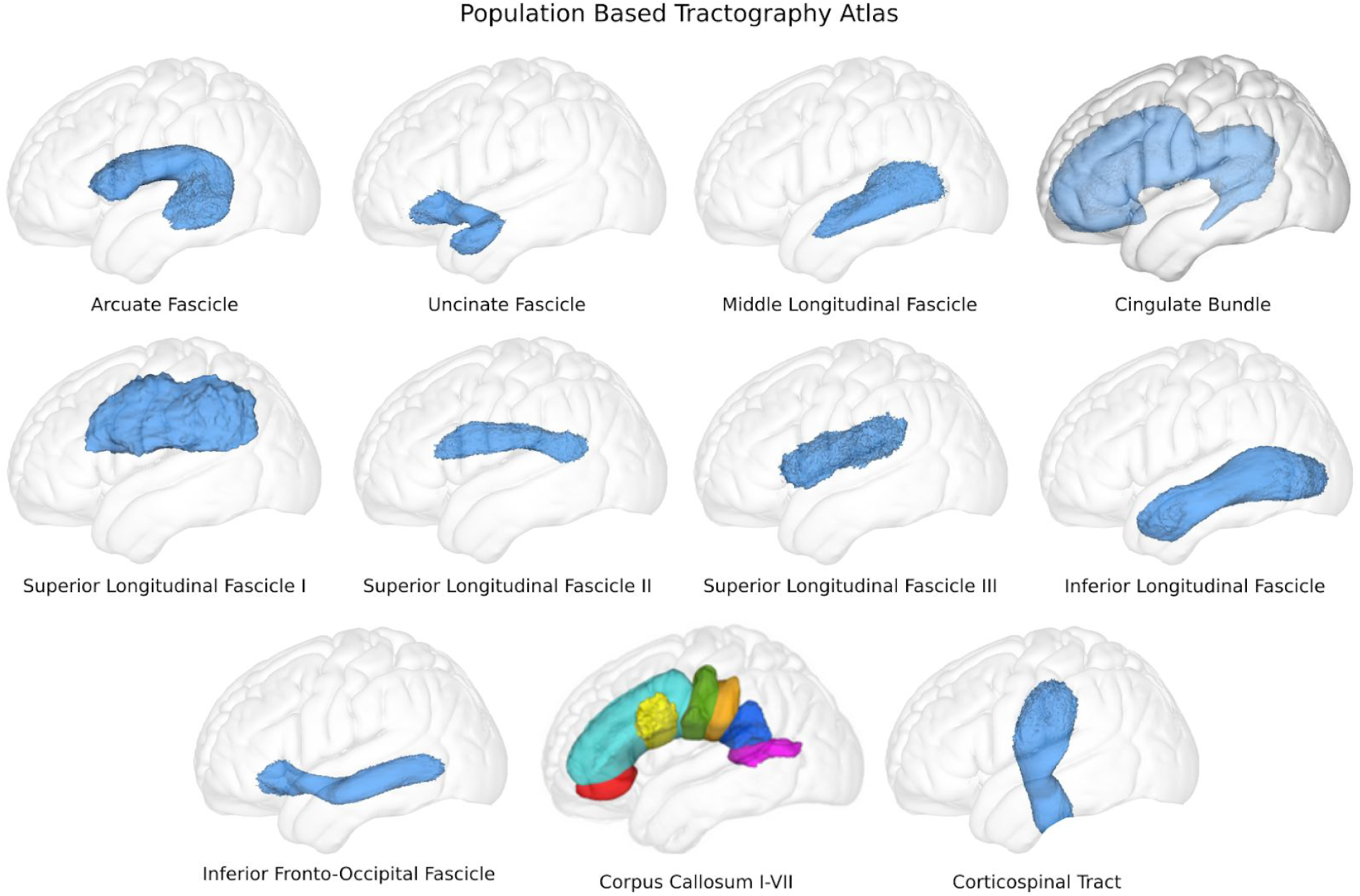
Population based tractography atlas. Obtained by aggregating the binary visitation maps of 600 subjects, and rendering the isosurface of the core of the tracts by thresholding them at the 10% level.

### 3.5. Population Maps of Cortical Connectivity

To assess the cortico-cortical connectivity at the population level, similarly to the volume based tract map, we computed a binary connectivity map on the cortical surface of each subject. We labeled each vertex of the surface if a streamline passed less than one voxel away from it, in the native space of the diffusion images. By doing so, we take into account every point on the cortex that could potentially be connected to the tract, thus reducing false negative connectivity results. Then, leveraging that the cortical surfaces are coregistered across subjects, we aggregated the individual binary maps to a group population map. The resulting map once more yields a natural probabilistic map of cortico-cortical connections of the different tracts. In this case, each vertex value denotes in how many subjects the vertex is reached by the tract.

Figure 5 shows the resulting connectivity maps projected in the average inflated surface, computed from 400 HCP subjects (Glasser *et al*., 2013). The maps are thresholded to the 5% level.

**Figure 5.**
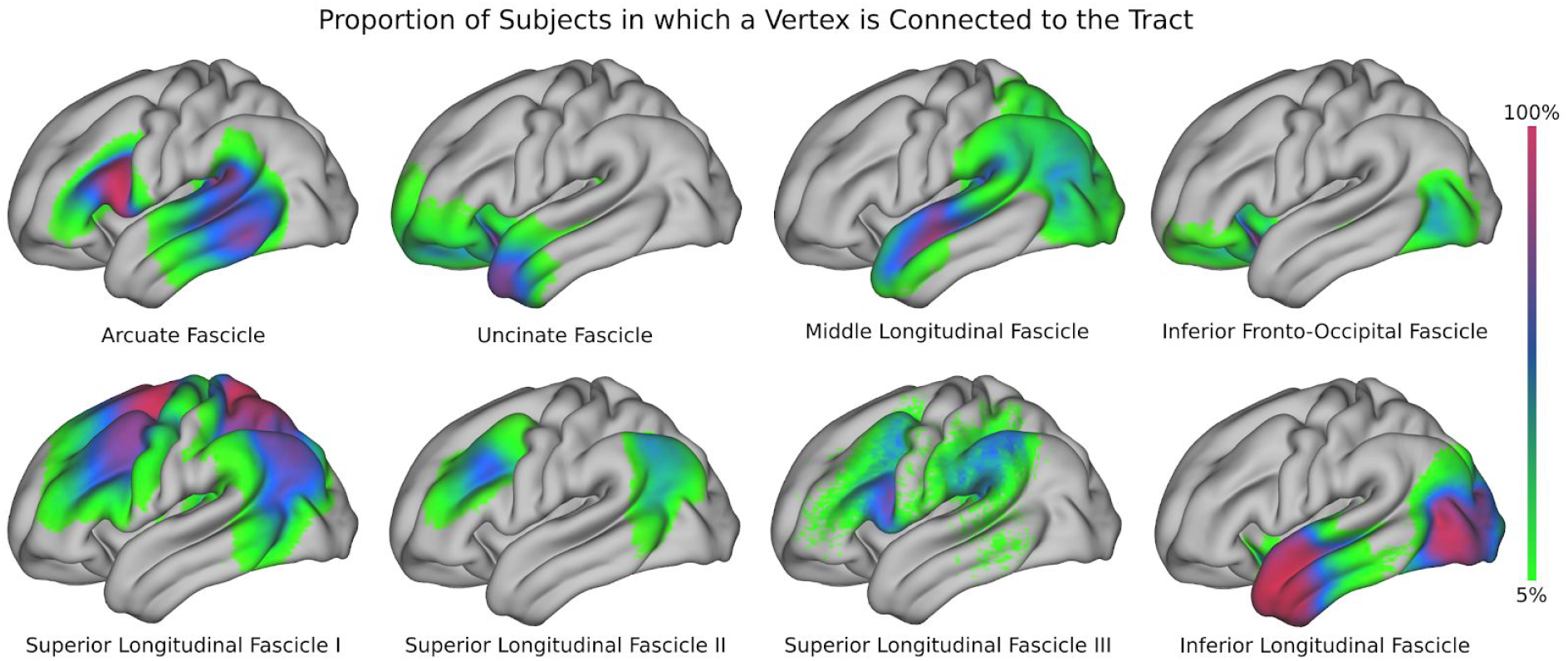
Population based connectivity maps of exemplary pathways. The colors represent in which percentage of subjects each tract connects to a vertex.

### 3.6. Assessing Lateralization of Language Pathways

To evaluate the sensitivity and robustness of our automatic tractography algorithm, we carried out a study of lateralization of the language pathways. Currently, there exist at least 3 main measures used to compute lateralization of pathways based on streamline tractography. If *S_h_* is the number of streamlines on one of the hemispheres (*h* ∈ [*L, R*]), and *v_h,i_* is the number of streamlines visiting voxel i in hemisphere h, the indices can be expressed as:

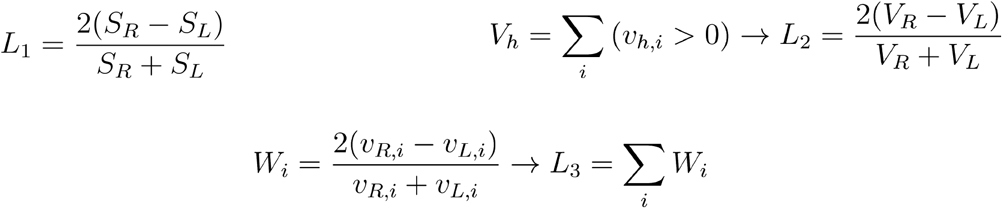

*L*_1_ directly compares the number of streamlines generated for the tract on each hemisphere (Thiebaut de Schotten et al., 2011). *L*_2_ compares the volume of the tract in each hemisphere (computed as the number of voxels visited by at least one streamline) (Thiebaut de Schotten et al., 2011). Finally, *L*_3_ compares the number of visited voxels, but weighted by the amount of streamlines passing through them (Matsuo et al., 2012). The 3 indices return a number between −2 and 2, where 0 indicates no lateralization, and the extremes −2 and 2 represent total left (−2) or right (2) lateralization respectively.

We computed the 3 lateralization indices for the main tracts associated with the language system: AF, SLF2, SLF3, IFOF, ILF, and UF (Gierhan, 2013). The resulting indices were highly correlated (r > 0.9 on every tract, Table 1), for which we decided to only use *L*_1_ for presenting results.

**Table 1.** Correlation between lateralization indices across subjects

|  | AF | IFOF | ILF | UF | SLF2 | SLF3 |
| --- | --- | --- | --- | --- | --- | --- |
| $L_1$ vs. $L_2$ | 0.96 | 0.98 | 0.90 | 0.92 | 0.99 | 0.99 |
| $L_1$ vs. $L_3$ | 0.99 | 0.99 | 0.99 | 0.97 | 0.99 | 0.99 |
| $L_2$ vs. $L_3$ | 0.96 | 0.98 | 0.90 | 0.97 | 0.99 | 0.99 |

Figure 6a presents the distributions obtained for the lateralization of each tract. The resulting distributions are consistent with previous studies: the AF and ILF are highly left lateralized (Propper *et al*., 2010; Thiebaut de Schotten *et al*., 2011; Gierhan, 2013; Panesar *et al*., 2018; Warrington *et al*., 2020); the SLF II, SLF III, and UF are highly right lateralized (Thiebaut de Schotten *et al*., 2011; Howells *et al*., 2018; Warrington *et al*., 2020), and the IFOF is slightly right lateralized (Thiebaut de Schotten *et al*., 2011; Wu *et al*., 2016; Warrington *et al*., 2020). To help interpreting the lateralization index, we further computed the lateralization ratio from *L*_1_. This ratio between the streamline count of the left and right hemisphere is a more intuitive measure of laterality, and allows us to easily interpret a value of *L*_1_ = 0.4 as “1.5 times more streamlines on the right hemisphere”. Furthermore, using a cumulative histogram, we can easily interpret the degrees of lateralization of each tract. For example, we can see that in the AF 70% of the population has at least 1.5 times more streamlines in the left hemisphere, as shown in the cumulative histograms of figure 6b.

**Figure 6.**
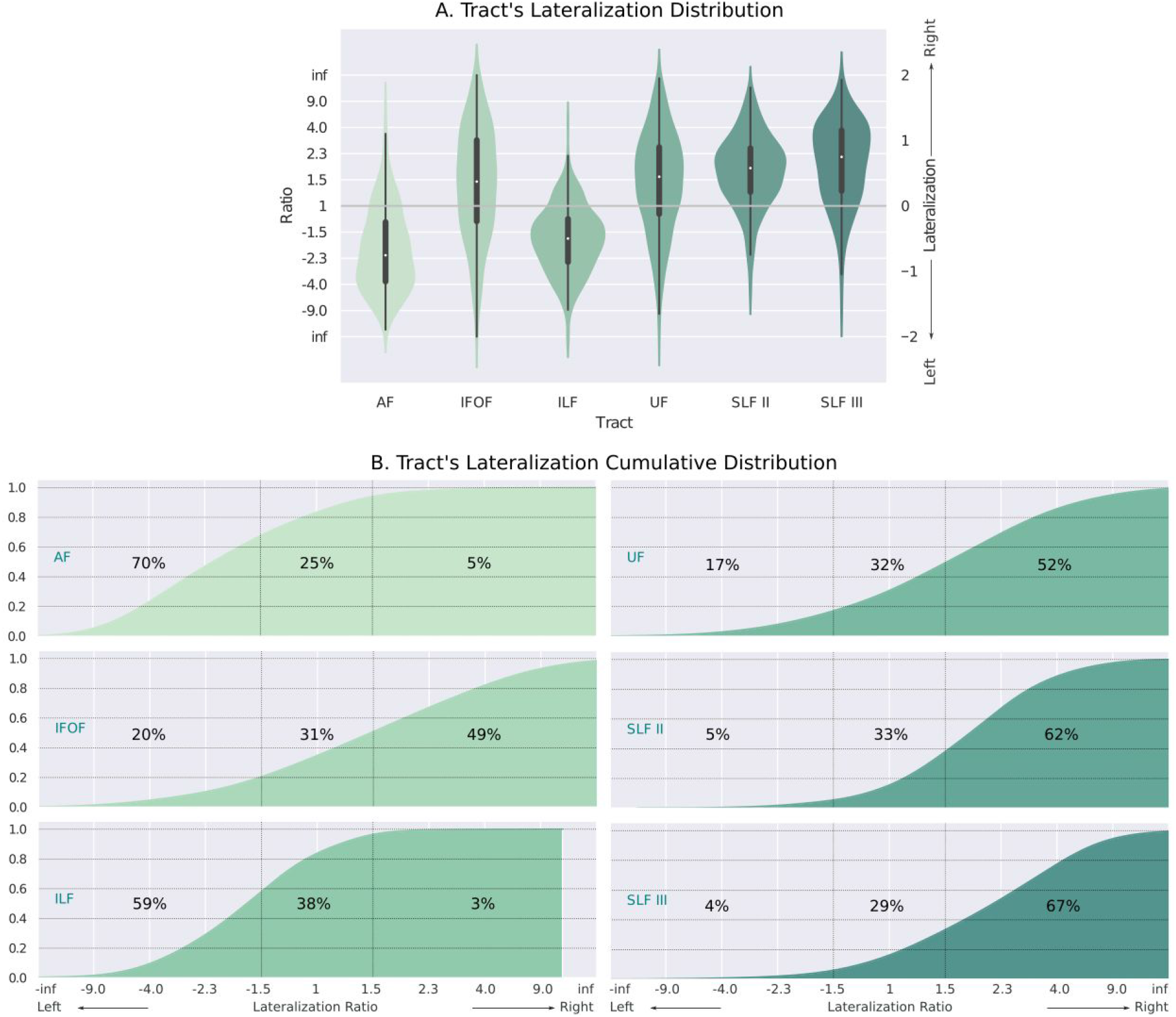
A. Distribution across subjects of the lateralization ratio for major language-related pathways. B. Cumulative distributions per tract. We labeled the percentage of subjects with 1.5 times more streamlines in the left (leftmost number) and right (rightmost number) hemisphere

### 3.7. The Role of Gender and Handedness in Lateralization

It has been reported in the literature that lateralization can be driven by handedness or gender of the subject (Thiebaut de Schotten *et al*., 2011; Howells *et al*., 2018). We now leverage our method to test if those individual differences are confirmed by the automatically computed lateralization of language-related tracts. We first divided our sample into left handed and right handed subjects using the handedness score, available in the HCP dataset. The score ranges from −100 to 100, where negative values indicate left-handedness, and positive ones right-handedness (Schachter et al., 1987). In this comparison, we included left-handed subjects with a score lower than −50, and right-handed subjects with a score higher than 50. The resulting groups had 75 left-handed individuals, and 525 right-handed ones. Given that lateralization values are not normally distributed, we compared them across groups using a non parametric U-test.

Figure 7 shows the lateralization values obtained when dividing the population based on handedness. The U-test revealed no relationship between handedness and lateralization (all p-values > 0.3).

**Figure 7.**
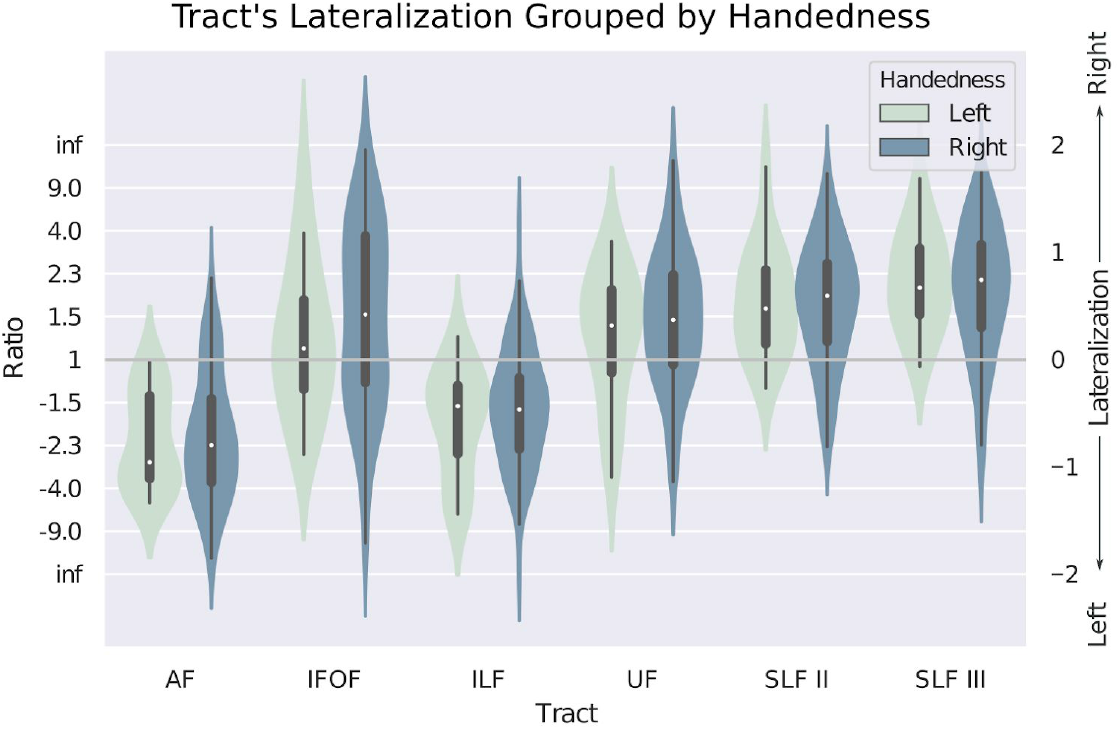
Distribution of the lateralization of each tract grouped by handedness. We found no statistically-significant difference between groups.

Finally, to evaluate previous studies on gender differences in lateralization of the language pathways in a large cohort, we divided our original sample into male and female subjects. Once again, we compared the lateralization in both groups using a non parametric U-test. Figure 8a shows the lateralization values obtained for each group for different tracts. In this case, and as shown in Table 2, only the IFOF (p<0.001, U=35405) and the ILF (p<0.03, U=10693) showed a significantly different distribution of lateralization values. However, it is important to remark that, while the difference is statistically significant, the cumulative distributions (Figure 8b) show that in both populations the overall lateralization trend is not changed. If we look at the lateralization of the IFOF, in both genders the IFOF is still mostly right lateralized, with 54% of females, and 44% of males having a ratio > 1.5. This is, 54% of females and 44% of males have 1.5 times more streamlines computed in the left hemisphere than in the right one. Meanwhile, the ILF is still highly left lateralized in both genders, with 55% of females, and 65% of males having a ratio < −1.5.

**Figure 8.**
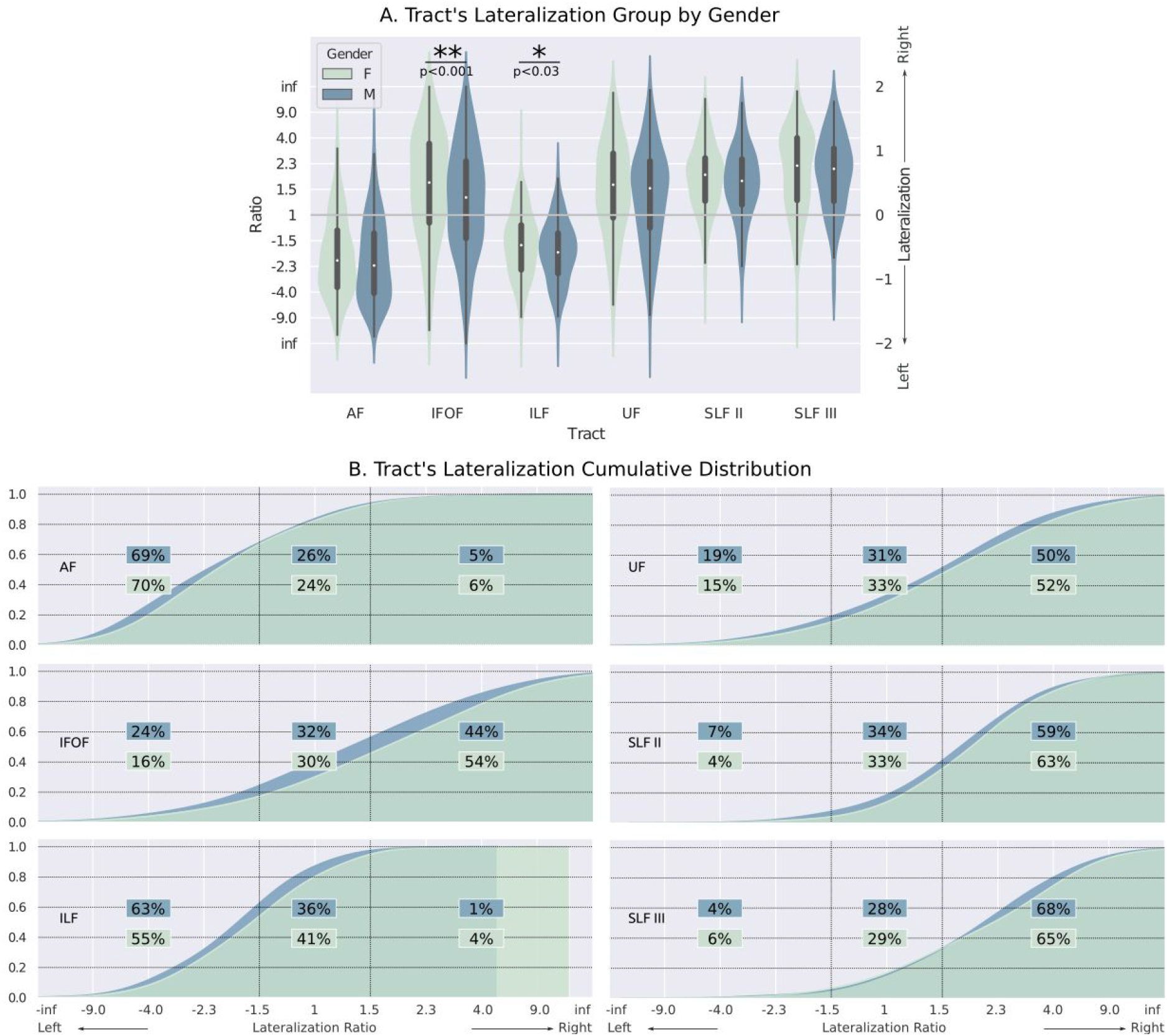
A. Distribution of the lateralization (and ratio) grouped by gender, only the IFOF and ILF showed significant differences (see table 2). B. Cumulative distributions per tract. We labeled the percentage of subjects with 1.5 times more streamlines in the left (leftmost number) and right (rightmost number) hemisphere.

**Table 2.** Results of U-test when comparing lateralization grouped by gender

|  | AF | IFOF | ILF | UF | SLF2 | SLF3 |
| --- | --- | --- | --- | --- | --- | --- |
| U-statistic | 44995 | 35405 | 10693 | 22948 | 22651 | 5733 |
| p-value | 0.06 | <0.001 | <0.03 | 0.07 | 0.07 | 0.26 |

**Table 3:**
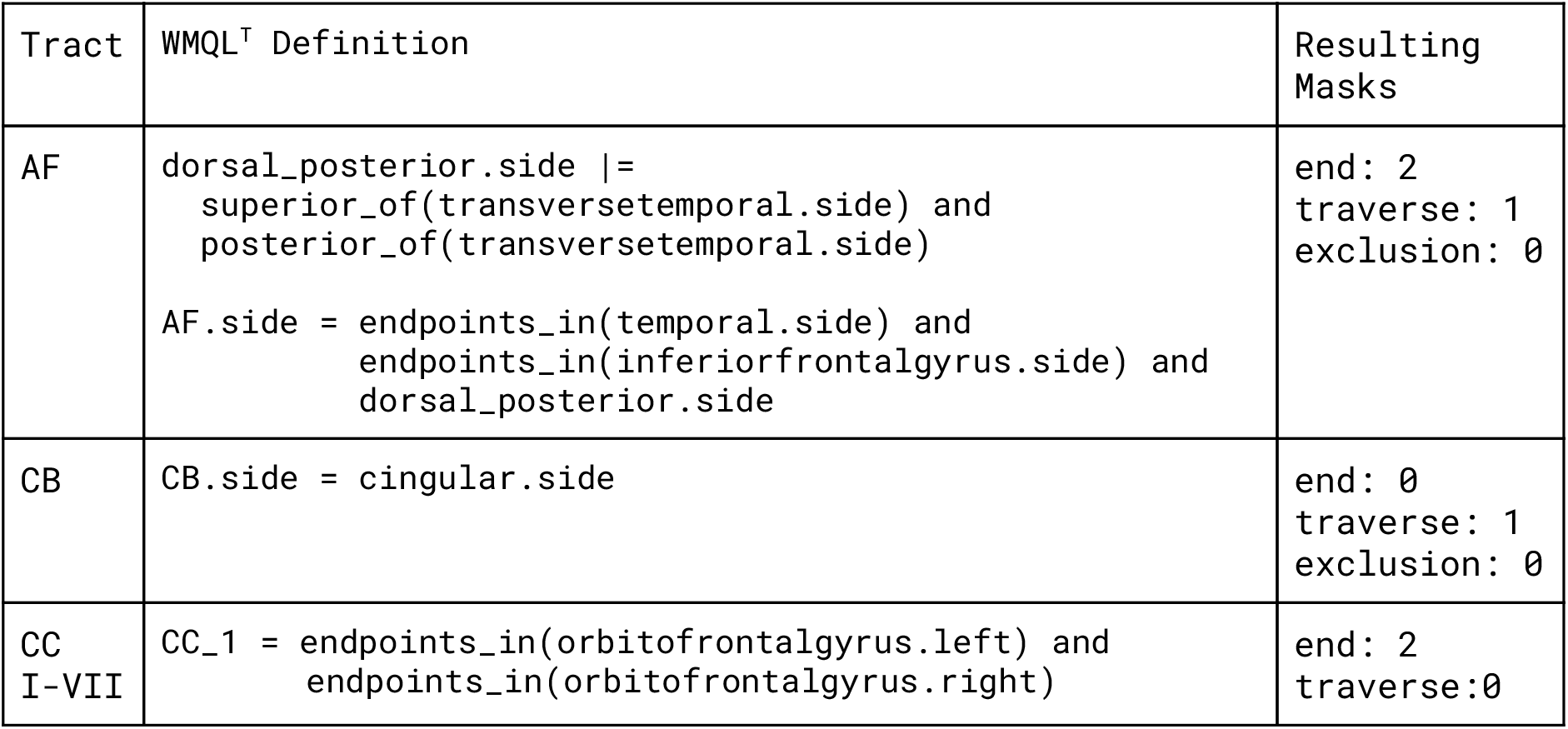
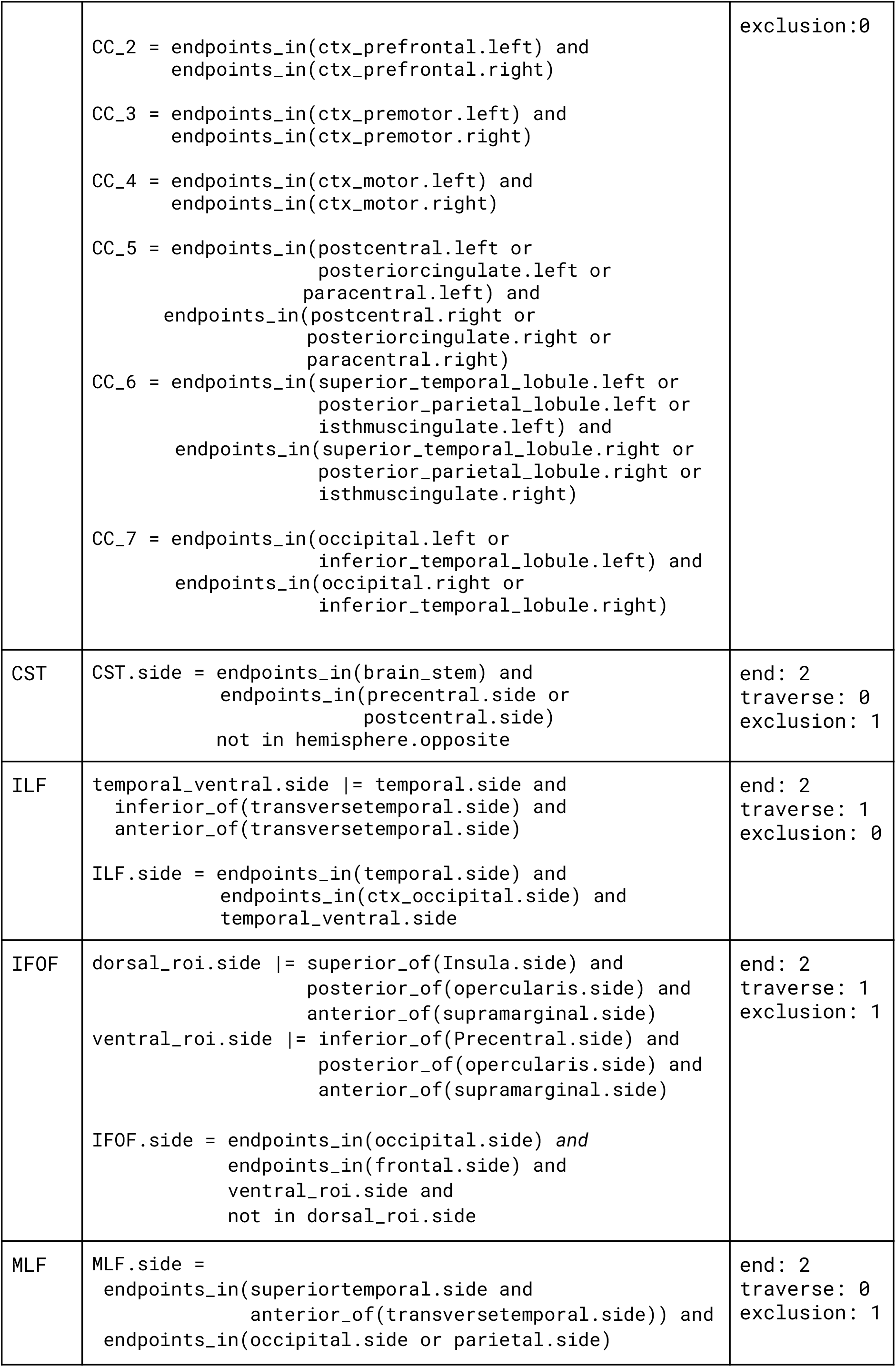
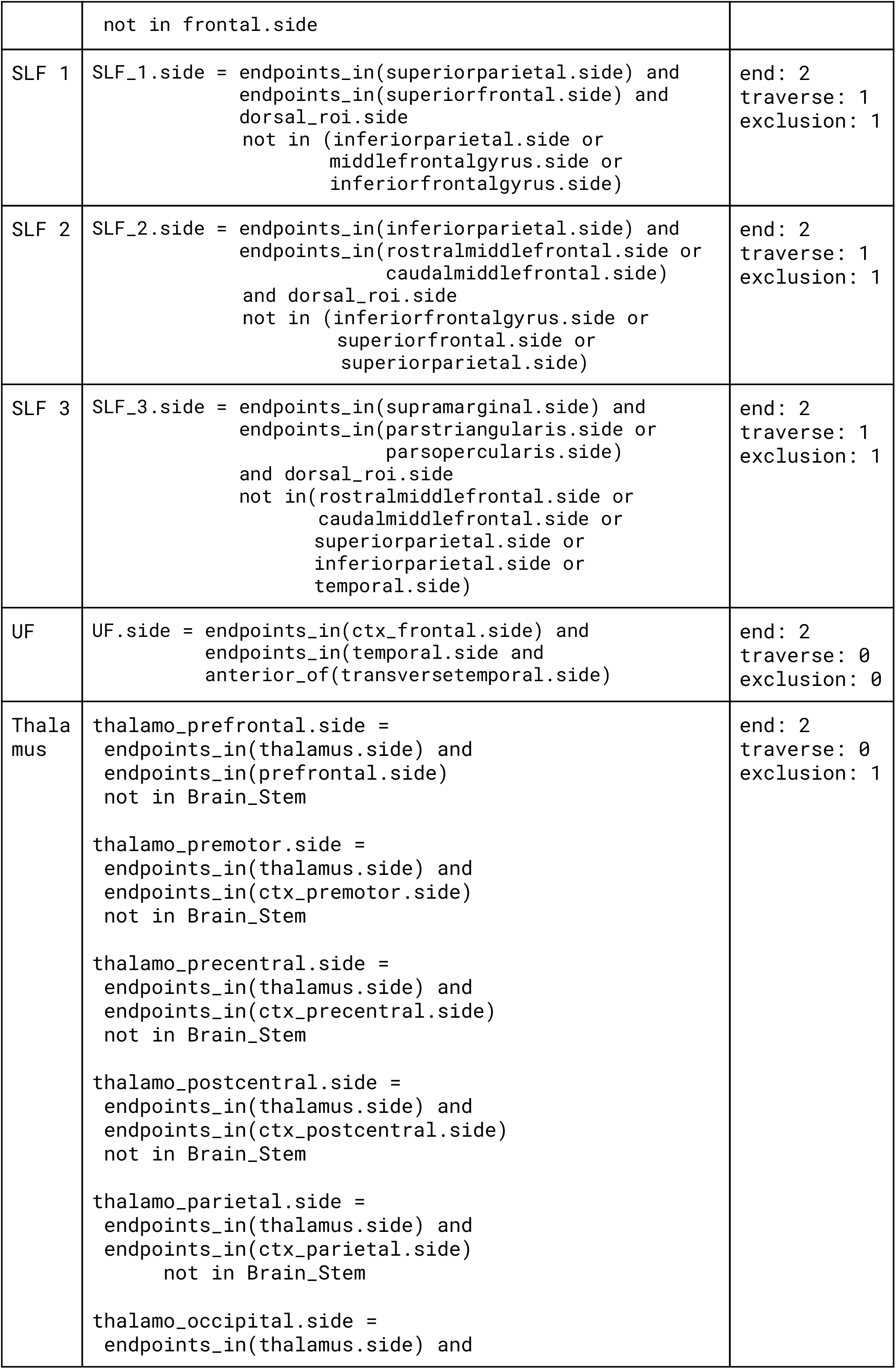
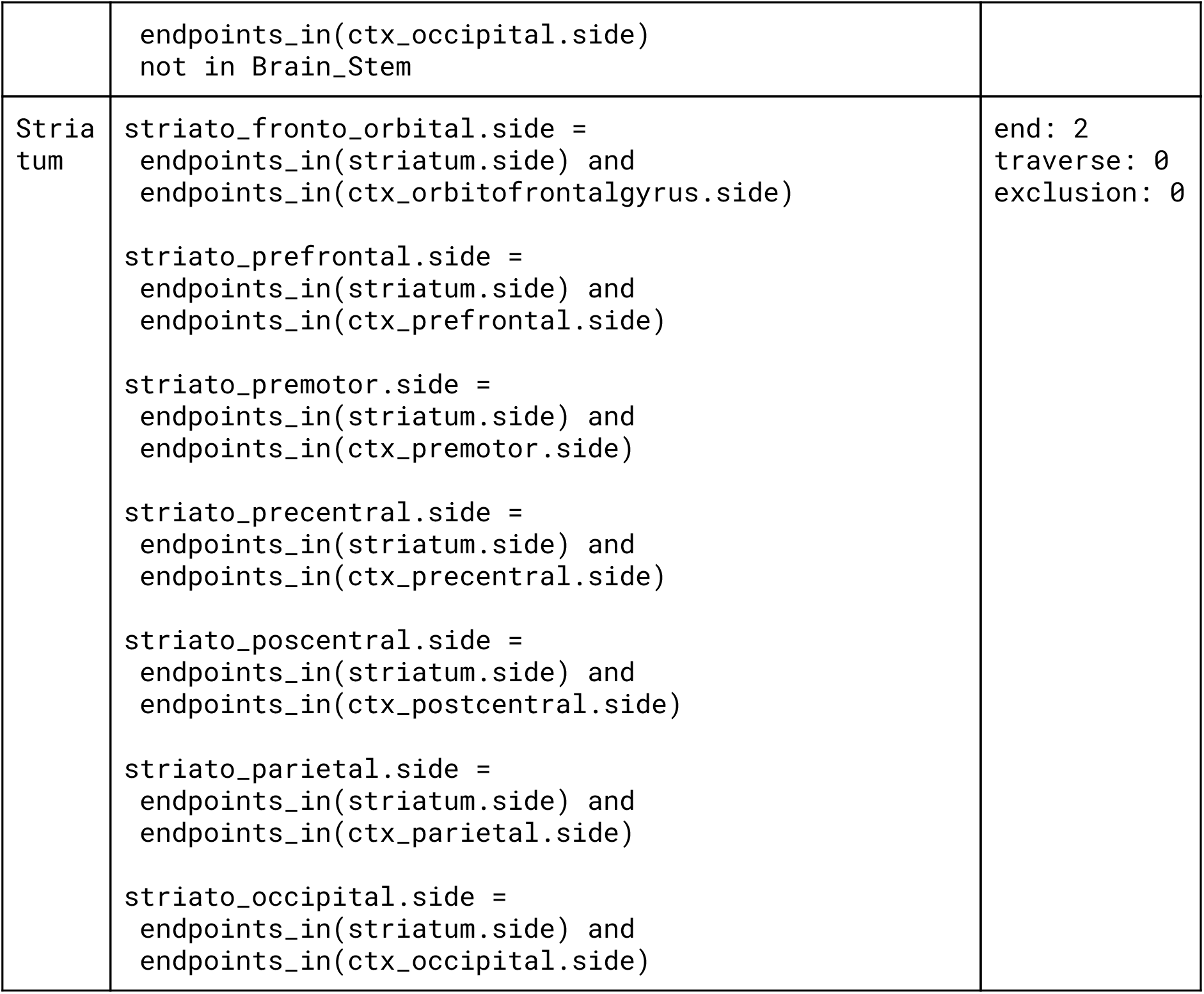
Definitions for 19 tracts and their subdivisions: arcuate fascicle (AF); cingulum bundle (CB); corpus callosum (I-VII); cortico-spinal tract (CST); inferior fronto occipital fascicle (IFOF); inferior longitudinal fascicle (ILF); middle longitudinal fascicle (MLF); superior longitudinal fascicle (SLF I, II, III); uncinate fascicle (UF); and connections to the thalamus and striatum. Based on the WMQL definitions by Wasserman et al. (2016)

## 4. Discussion

In this work we introduced a method that reconstructs major brain fiber pathways from their textual descriptions. This allows an objective and transparent definition of white matter pathways for group analysis of tractography, while being flexible to define new tracts for a given study. Furthermore, it can be used to create tracking masks compatible with current ROI-based reconstruction methods, such as AFQ or XTRACT, enabling to easily extend them beyond their predefined set of tracts.

In order to define brain pathways, our method builds on top of the existing White Matter Query Language. Particularly, we define a more constrained and tractography-oriented language, namely WMQL^T^. As shown in section 2.1, WMQL^T^ syntax is easy to interpret, and its expressiveness allows representing major white-matter tracts. This makes it easy for the user to extend our predefined set of 19 tracts, overcoming thus one of the major limitations of current reconstruction methods (Zhang *et al*., 2008; Yendiki *et al*., 2011; Yeatman *et al*., 2012; Warrington *et al*., 2020). It is important to remark that WMQL^T^ definitions rely on atlas regions, for which the spatial location and granularity of the definitions is constrained by the chosen atlas. However, since any atlas can be used (i.e. structural or functional ones), WMQL^T^ is flexible enough to allow defining a same tract using multiple atlases, therefore resulting in different levels of granularity. Moreover, WMQL^T^ enables to define tracts using multiple atlases from different modalities, as long as they have non-overlapping labels. This allows, for example, to define labels aggregating anatomical and functional information.

The current WMQL’s interpreter translates language terms into rules for filtering a whole-brain tractography (Wassermann *et al*., 2016). Because of this, reconstructing a specific tract becomes a time-consuming and inefficient process, needing to track millions of streamlines in order to filter a few representatives. In contrast, our WMQL^T^ interpreter (section 2.2) translates WMQL queries into tracking masks. By constraining the tracking to those masks, it is possible to drastically reduce the time needed to reconstruct a specific tract, while still simulating a large number of streamlines per seedpoint. More importantly, our tracking masks can be used in any tractography software (i.e. MRtrix, FSL, Dipy), giving the user freedom to use their preferred implementation. This further allows us to complement existing reconstruction methods. For example, masks created on the MNI or F99 templates can be used to easily extend XTRACT (Warrington *et al*., 2020) and leverage their tracking techniques.

To ease the process of reconstructing subject-specific tracts, we implemented a tractography pipeline (section 2.3), allowing us to easily translate WMQL^T^ definitions into streamlines. Our pipeline leverages state-of-the-art diffusion models (Tournier *et al*., 2007) and tracking algorithms (Tournier *et al*. 2012). We seed from both *transverse* and *end* masks in order to better characterize both the core and ending points of each tract (Warrington *et al*., 2020).

Our Technique Makes it Easier to Translate Neuroanatomical Knowledge into Tracts Using our pipeline, we reconstructed 19 tracts in 600 subjects from the Human Connectome Project. Both the single-subject and population-wise results show that the morphology (Figure 2 and 4) and connectivity (Figure 5) of the reconstructed tracts is in accordance with descriptions from the current literature (O’Donnell and Westin, 2007; Catani and Thiebaut de Schotten, 2008; Yeatman *et al*., 2012; Wassermann *et al*., 2016; Warrington *et al*., 2020). Furthermore, our results highlight the potential of our tool to study commonalities and individual variations in a population. By combining the resulting tracts and visitation maps in a common space, we naturally derived probabilistic population maps, representing the probability of finding a tract in a specific position.

### WMQL^T^ Allows to Easily Define and Reconstruct Tracs using Multiple Atlases

Our WMQL^T^ interpreter can handle multiple atlases concurrently, allowing us to write richer pathway definitions. We showcased this by defining and reconstructing 3 functional subcomponents of the corticospinal tract (Figure 3). This capability of our method enables us to easily define and systematically reconstruct brain pathways based on multimodal information.

### Lateralization Indices are Highly Correlated in Language-Related Pathways

Currently there is no gold standard on how to quantify lateralization from dMRI tractography. Having reconstructed each tract as a set of streamlines allowed us to compute and compare the 3 most common ways to assess lateralization: counting streamlines, counting voxels, and their weighted combination (section 3.6). Our results showed that the 3 indices are highly correlated in language-related tracts (Table 1). This helps to unify results obtained across papers using different lateralization indices.

### Evaluation of the Lateralization of Language-Related Pathways

To further validate our technique, we computed the lateralization of 5 language-related tracts: AF, IFOF, ILF, UF, SLF2, and SLF3. The obtained distributions closely follow neuroanatomical knowledge, as well as results reported by another work (Warrington *et al*., 2020). In order to better understand the indices obtained for lateralization, we transformed them into a ratio, expressing how many times more streamlines are computed in one hemisphere. Our results show that AF is highly left lateralized, as reported in the literature (Thiebaut de Schotten *et al*., 2011; Gierhan, 2013), with 70% of the population having 1.5 times more streamlines on the left side. The ILF is also left lateralized, as shown in the literature (Panesar *et al*., 2018), with 59% of the population showing a ratio < −1.5. The IFOF was right lateralized, as reported by (Warrington *et al*., 2020; Thiebaut de Schotten *et al*., 2011), with 49% of the population showing 1.5 more streamlines in the right than left hemisphere. However, it is important to notice that the cumulative distribution (Figure 6b) shows that 20% of the population is highly left lateralized. This underlines the benefit of translating lateralization indices into the more intuitive lateralization ratio. Finally, UF, SLF2 and SLF3 were right lateralized, as reported by (Thiebaut de Schotten *et al*., 2011). In particular, the IFOF showed in 49% of the population a ratio > 1.5, while the UF, SLF2, and SLF3 in 51%, 62% and 67% respectively. Our resulting lateralization, and its correspondence with current knowledge, highlight the ability of our technique to correctly reconstruct white matter tracts in large cohorts.

### Only the Inferior Fronto Occipital Fascicle and Uncinate Fascicle Showed Gender Differences in Lateralization

We found no significant differences in lateralization when grouping our subjects by handedness, and only the IFOF and UF showed significant group differences when dividing by gender. However, this did not translate to a change in the general trend of lateralization. Respect to the IFOF, both groups were left lateralized, with 54% of females and 44% of males showing a ratio > 1.5. This means that 54% of females, and 44% of males have 1.5 times more left streamlines than right ones. Meanwhile, the ILF is also highly left lateralized in both groups, with 55% of females, and 63% of males having a ratio < −1.5. Such results highlight the advantage of computing ratios and looking at the general distribution of lateralizations, as well as the potential of our technique to study pathway properties in large populations.

## 5. Conclusion

Translating neuroanatomical knowledge into methods for tract reconstruction remained challenging for the non-technical users. Recent advances in automated reconstruction methods allowed to isolate major tracts in large populations, but remained difficult to expand with new definitions. In this work, we presented a method to define and reconstruct white-matter tracts using a near-to-english language. Our technique can be used by itself or complementing existing ones to automatically reconstruct major tracts in large populations.

By defining and reconstructing major brain tracts, we showed how our method consistently creates tracts in accordance with the neuroanatomy literature. This was further validated by comparing the lateralization of five language-related tracts. We also showcased two applications of our tool: how to enrich tract definitions by using multimodal information, and how to assess group differences in tract-specific properties.

Both our tractography pipeline, as well as the results will be available on GitHub. We expect our tool to help to improve our knowledge on white-matter organization, and its variability across subjects. Future iterations on the language will further improve its expressiveness power, therefore increasing the capabilities of our pipeline.

## 6. Acknowledgments

This work has received funding from the ERC-StG NeuroLang.

## Notes

### Competing Interest Statement

The authors have declared no competing interest.

